# A TMEM175 Mutation Disrupts the Lysosomal-Mitochondrial Axis Causing Short QT Syndrome and Sudden Cardiac Death

**DOI:** 10.64898/2026.11.05.764745

**Authors:** Juan Manuel Ruiz-Robles, Isabel Martínez-Carrascoso, Álvaro Macías, Ana I. Moreno-Manuel, Patricia Sánchez-Pérez, Marre E. Corver, Paula G. Socuéllamos, Amaia Talavera-Gutiérrez, Francisco J. Bermúdez-Jiménez, Eva Cabrera-Borrego, Rosa Macías-Ruiz, Fernando Martínez de Benito, Juan Pablo Ochoa, Sean Jurgens, Connie R. Bezzina, Juan Jiménez-Jaimez, José Jalife, Francisco M. Cruz

## Abstract

Short QT Syndrome (SQTS) is a rare cardiac channelopathy characterized by abnormally short QTc intervals and increased sudden cardiac death (SCD) risk. Only 20-30% of SQTS cases have been linked to genetic mutations, typically in surface membrane ion channels. We identified a novel lysosomal *TMEM175* gene mutation (p.Arg377His) in 7 patients in a family with SQTS and SCD. Genetic analyses, including whole exome sequencing in the family and rare-variant burden testing in ∼118,000 UK Biobank participants, suggested an association between *TMEM175* rare variants and QT interval shortening. Functional studies in CRISPR-engineered mice and patient-specific induced pluripotent stem cellderived cardiomyocytes (iPSC-CMs) revealed that TMEM175^R377H^ causes shortened QT intervals and hastened repolarization, respectively, as well an increased arrhythmia susceptibility. Isolated TMEM175^R377H^ cardiomyocytes showed action potential duration (APD) abbreviation, confirmed in whole TMEM175^R377H^ mouse hearts and patient-specific iPSC-CM monolayers. In silico and electron microscopy studies showed that TMEM175^R377H^ disrupts lysosomal-mitochondrial interactions, leading to mitochondrial dysfunction, as evidenced by the reduced maximal respiration and spare respiratory capacity, with reduced ATP production, I_KATP_ activation, and APD shortening. This mutation represents the first non-sarcolemmal ion channel defect associated with SQTS and high SCD risk, potentially expanding the genetic landscape of the disease.

## Introduction

Short QT Syndrome (SQTS) is a rare inheritable cardiac channelopathy characterized by QT interval shortening, predisposing to malignant ventricular arrhythmias and sudden cardiac death (SCD) in patients with structurally normal hearts^1^. The main ECG parameter for diagnosing SQTS is a corrected QT (QTc) interval consistently under 320 ms, without secondary causes, establishing a probable diagnosis, even without other symptoms^2^. Diagnosis should also be considered for QTc intervals between 320 and 360 ms, including a history of ventricular fibrillation without structural heart disease, family history of SQTS, unexplained SCD at ≤40 years, or a confirmed pathogenic mutation^3^. Fewer than 250 cases of congenital SQTS have been reported, with only 20-30% of patients with positive genetic tests, and four single genes associated with SQTS^4^. Highthroughput sequencing techniques such as Whole Genome Sequencing (WGS) and Whole Exome Sequencing (WES) have recently emerged as an avenue towards finding new causative genes in families affected by rare disorders, alongside conventional linkage techniques, and may similarly help identify novel causative genes in SQTS families.

TMEM proteins are a diverse family with distinct roles and pathophysiological mechanisms. For example, TMEM43, which is associated with the nuclear envelope, has been implicated in arrhythmogenic right ventricular cardiomyopathy (ARVC)^5^, while TMEM175 is a lysosomal protein previously associated with Parkinson’s disease through common variation according to GWAS^6^. The TMEM175 channel is known to regulate lysosomal pH, allowing proper progress of autophagy and regulating mitochondrial activity^7–10^. At the molecular level it acts through proton leak in the lysosome; i.e., as a counterpart of the V-ATPase responsible for lysosomal acidification^7^. Dysregulation of such an orchestrated process causes alterations in the flow of autophagy. Currently, Parkinson’s disease (PD) is the only condition associated with *TMEM175* variants, which act as a risk factor for PD rather than a monogenic cause, although there is no consensus on the specific pathogenic mechanism^6^. On the other hand, the relationship between TMEM175 and the regulation of mitochondrial activity has not been clearly demonstrated yet, even though physical lysosome-mitochondria interaction has been widely described in cardiomyocytes^11^. Thus, the TMEM175 channel might have an important yet undescribed role in the regulation of mitochondrial activity in cardiomyocytes. However, it is unknown whether the channel controls the cardiac action potential duration (APD).

In this work, we report on a newly identified SQTS family carrying the TMEM175^R377H^ mutation (GRCh38 chr4:958111G>A; c.1130G>A; p.Arg377His).

Genetic analyses, including WES with quartet analysis in a family affected by SQTS and rare-variant burden testing in ∼118,000 UK Biobank participants, suggested an association between *TMEM175* variants and QT shortening. In addition, our findings in a CRISPR-mediated gene-edited murine model and patientspecific induced pluripotent stem cell-derived cardiomyocytes (iPSC-CMs) support the causality of this variant for SQTS in the family. Specifically, these models reveal a novel mechanism mediated by the dysregulation of the cardiomyocytés energy state, which decreases intracellular ATP, activates KATP channels and shortens the APD and the QT interval. To our knowledge, this is the first mutation in a non-sarcolemmal ion channel associated with SQTS and a high incidence of SCD.

## Results

### Description of a new family with SQTS and SCD

We report a family profoundly affected by SCD. The proband (III.9; black arrow in **Figure 1A**) is a 16-year-old male referred for cardiac evaluation following two cases of familial SCD. His ECG showed a short QTc interval (320 ms), a Tpeak-Tend interval of 80 ms, and abnormal T wave morphology (**Figure 1B and Table 1**). He was asymptomatic, and cardiac imaging demonstrated a structurally normal heart. Both ambulatory ECG Holter monitoring and treadmill testing were unremarkable. Due to the significant family history of SCD, he was advised to receive an implantable cardioverter-defibrillator (ICD), which he declined. He began quinidine therapy at 1200 mg/day, but this resulted in no significant QTc interval prolongation. Holter monitoring revealed reduced shallow slope in the QTheart rate relationship compared to healthy and unrelated individual (**Supplemental Figure 1**), even during quinidine treatment, as expected from SQTS patients^12^.

**Figure 1.**
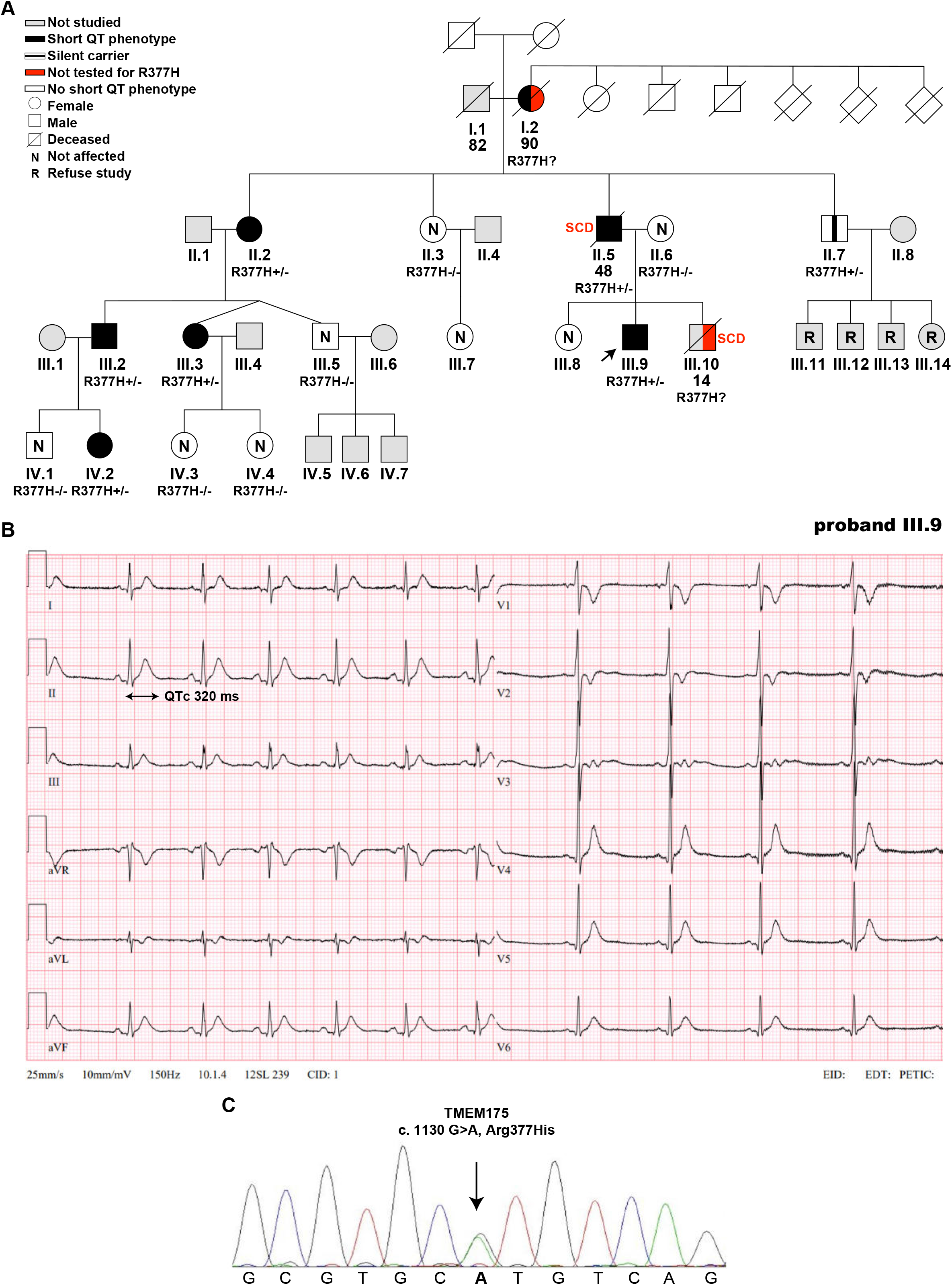
Arrhythmias and SCD in a family affected by the pathogenic TMEM175^R377H^ variant. **A)** Family pedigree according to the carrier status of the p.Arg377His *TMEM175* gene variant. The proband is marked with an arrow. Males are represented as squares and women as circles. Deceased patients are marked with a slash (/). Genotyped carriers are in black; likely carriers not genotyped are in red. Age at diagnosis is shown, as well as Sudden Cardiac Death events (SCD) and wild-type (WT) patients. **B)** Representative ECGs from individual III.9 proband exhibiting short QTc intervals (< 340 ms) and fulfilled current diagnostic criteria for the short QT syndrome. Scale: 1 mm/10 mV at 25 mm/sec. See Table 1 for numeric values. **C)** DNA Sanger sequencing chromatogram from proband’s (III.9) genomic DNA. The trace shows a heterozygous substitution of guanine to adenine resulting in the R377H (c.1130G>A (p. R377H)) amino acid change.

**Table 1.**
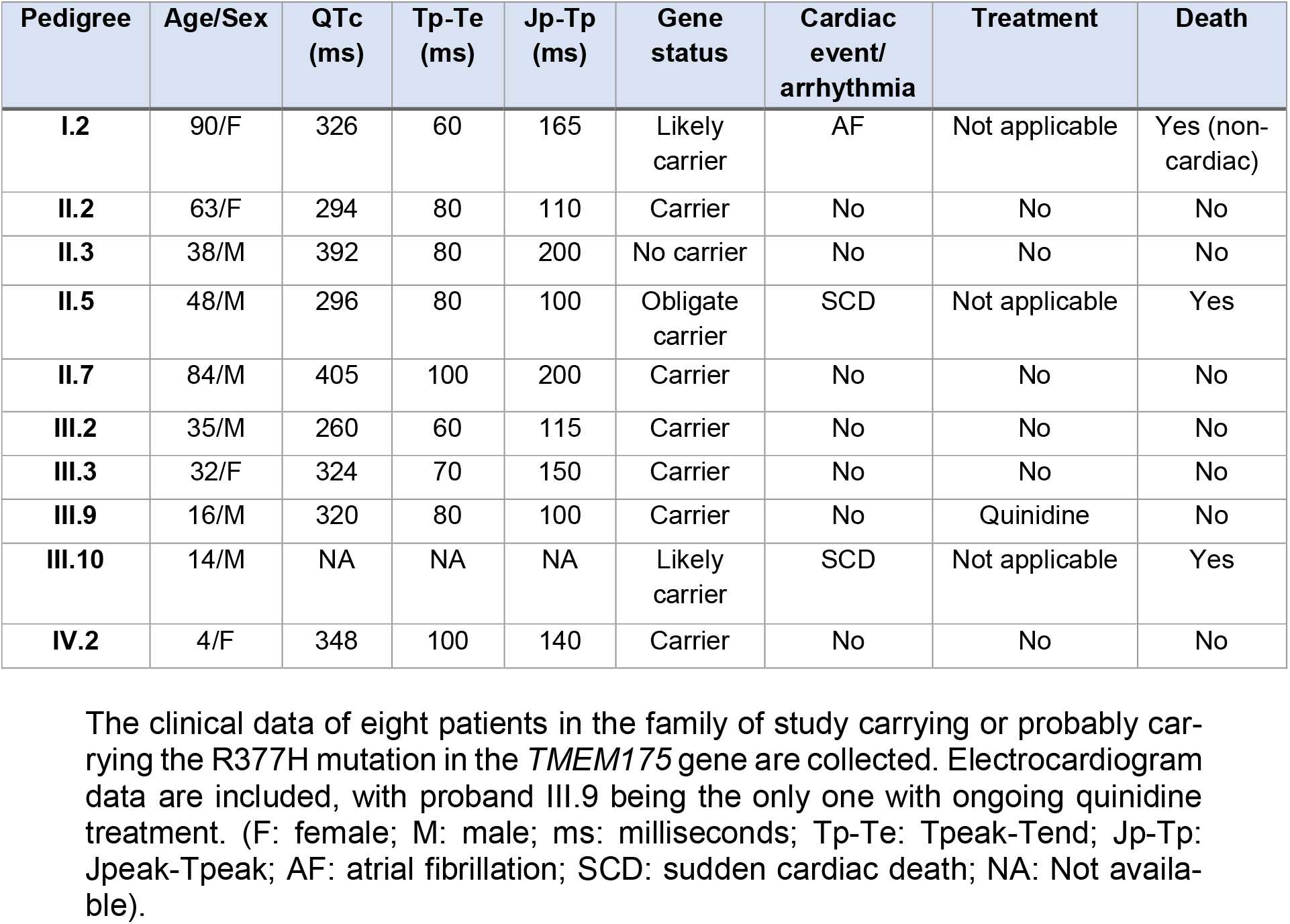
Clinical and genetic spectra of patients carrying the TMEM175^R377H^ mutation.

| **Pedigree** | **Age/Sex** | **QTc (ms)** | **Tp-Te (ms)** | **Jp-Tp (ms)** | **Gene status** | **Cardiac event/ arrhythmia** | **Treatment** | **Death** |
|---|---|---|---|---|---|---|---|---|
| **I.2** | 90/F | 326 | 60 | 165 | Likely carrier | AF | Not applicable | Yes (non-cardiac) |
| **II.2** | 63/F | 294 | 80 | 110 | Carrier | No | No | No |
| **II.3** | 38/M | 392 | 80 | 200 | No carrier | No | No | No |
| **II.5** | 48/M | 296 | 80 | 100 | Obligate carrier | SCD | Not applicable | Yes |
| **II.7** | 84/M | 405 | 100 | 200 | Carrier | No | No | No |
| **III.2** | 35/M | 260 | 60 | 115 | Carrier | No | No | No |
| **III.3** | 32/F | 324 | 70 | 150 | Carrier | No | No | No |
| **III.9** | 16/M | 320 | 80 | 100 | Carrier | No | Quinidine | No |
| **III.10** | 14/M | NA | NA | NA | Likely carrier | SCD | Not applicable | Yes |
| **IV.2** | 4/F | 348 | 100 | 140 | Carrier | No | No | No |

The clinical data of eight patients in the family of study carrying or probably car-rying the R377H mutation in the *TMEM175* gene are collected. Electrocardiogram data are included, with proband III.9 being the only one with ongoing quinidine treatment. (F: female; M: male; ms: milliseconds; Tp-Te: Tpeak-Tend; Jp-Tp: Jpeak-Tpeak; AF: atrial fibrillation; SCD: sudden cardiac death; NA: Not availa-ble).

After a 10-year follow-up, he remains asymptomatic. His brother (III.10) died suddenly at age 14 while at rest during the day, having never experienced syncope, participated in competitive sports, or undergone ECG evaluation. Postmortem examination was unremarkable, ruling out other causes and indicating a cardiac event as the cause of death. Eight years later, their father (II.5; **Figure 1A** and **Table 1**) died suddenly at age 48 while resting. When this occurred, the proband (III.9) was 6 years old; the father had never experienced syncope or other cardiac symptoms, however, a recovered ECG showed a reduced QTc interval of 296 ms (**Table 1 and Supplemental Figure 2-III**).

A clinical and ECG-based familial cascade screening identified 5 additional affected family members. 12-lead ECG recordings (**Supplemental Figure 2I-V**) and clinical phenotypes of affected family members are shown in **Table 1**. All had marked QT interval shortening but were asymptomatic and had no major cardiac events. Transthoracic echocardiograms, treadmill tests, and Holter recordings were unremarkable in all affected members, and none had been started on quinidine or had an ICD implanted. QTc intervals tended to be shorter in males than females (293±31 ms *vs* 323±22 ms, p=0.20). Additionally, the incidence of major arrhythmic events was higher in male family members, with 2 SCD events in males and none in female carriers. According to clinical data, no events of PD or related neurological syndromes have been detected in this family, including among elderly family members.

### Whole-exome sequencing identified a variant in *TMEM175* as the likely causal variant

An NGS panel including 380 genes related to cardiomyopathies (**Supplemental Table 1**), cardiac arrhythmia and SCD (including genes *KCNH2*, *KCNQ1*, *KCNJ2* and *SLC4A3* from the SQTS ClinGen panel) was tested in the proband, with no relevant variant detected based on standard clinical genetics assessments.

Without a molecular diagnosis, we proceeded to whole exome sequencing (WES) in a quartet analysis on three affected (II.2, III.3 and III.9) and one unaffected (II.3) family members, using an approach similar to previously described^13^ (see Materials & Methods for details). Consistent with the family pedigree, we retained only heterozygous variants present in the three affected members and absent in the unaffected sibling, reflecting an autosomal dominant pattern of inheritance (**Figure 1**). This initial quartet analysis resulted in 95 candidate variants. A list of the 95 initial variants and details are available in the Online Supplemental.

To pinpoint the causal variant, we applied a robust step-wise prioritization. First, synonymous variants without a presumed protein effect were excluded, reducing the list to 70 plausible candidates. Next, we applied a filtering based on SQTS population frequency, excluding variants with minor allele frequency (MAF) >0.01%, which further reduced the list to 18 candidate variants (**Table 2**). This threshold was selected to account for incomplete penetrance, a known feature in cardiac syndromes. From that list, none of the variants corresponded to genes previously described in SQTS. Two in-frame indel variants were also ruled out as they were located in a proline-rich region of variable length *(HTT)* or in a predicted disordered region of the protein *(ANKRD11*). Finally, we applied a composite score integrating eight i*n silico* predictors to assess missense variant effect (SIFT, PolyPhen-2, LRT, MutationTaster, MutationAssessor, FATHMM, PROVEAN, and MKL). This comprehensive approach identified a heterozygous missense mutation in the *TMEM175* gene (p.Arg377His; GRCh38 chr4:958111G>A; c.1130G>A) as the most likely causal variant with 7 out of 8 predictors indicating a deleterious effect (see **Table 2** and *Online Supplemental* for prioritization scoring), more than any other prioritized variant. Further details on variant filtering and prioritization scoring are described in *Supplementaly Material*.

**Table 2.** Prioritization according to protein effect prediction of candidate variants (MAF<0.01%). Pathogenicity scores and qualitative predictions of missense variants across multiple algorithms. In frame indels were not scored (NA) due to *in silico* tools limitations, and therefore ruled out from the downstream analysis. Abbreviations for predictions: D, Damaging / Disease causing; B, Benign; T, Tolerated; P, Possibly damaging; N, Neutral; M, Medium impact; L, Low impact; U, Unknown; NA / –, Not available or not applicable. The final “Score” represents the cumulative number of pathogenic predictions. Please, note rs751828848 in TMEM175 reached the highest score.

| **Symbol** | **SNP ID** | **SIFT** | **Polyphen2** | **LRT** | **MutationTaster** | **MutationAssessor** | **Fathmm** | **PROVEAN** | **MKL** | **Score** |
|---|---|---|---|---|---|---|---|---|---|---|
| TMEM175 | rs751828848 | **D** | **D** | **D** | **D** | **M** | T | **D** | **D** | **7** |
| ANKRD11 | rs757738828 | NA | NA | NA | NA | NA | NA | NA | NA | **NA** |
| HTT | rs547233813 | NA | NA | NA | NA | NA | NA | NA | NA | **NA** |
| AQR | rs763883158 | T | B | N | N | N | **D** | N | N | **1** |
| TMCO3 | rs41288604 | T | B | N | N | N | T | N | **D** | **1** |
| BIRC7 | rs781384192 | T | B | N | N | N | T | N | N | **0** |
| TBC1D2 | rs199544260 | T | B | N | N | N | T | N | N | **0** |
| SNTB2 | rs542537218 | T | B | N | N | N | T | N | N | **0** |
| TAS2R46 | rs768843374 | T | B | - | N | N | T | N | N | **0** |
| CLEC18C | rs3869428 | T | B | N | N | L | T | N | N | **0** |
| GTPBP8 | rs139947240 | T | **D** | N | **D** | **M** | T | N | **D** | **4** |
| NOP14 | rs149423815 | T | P | N | N | L | T | N | N | **0** |
| GRK4 | rs142548315 | T | B | U | **D** | L | T | N | **D** | **2** |
| NOP14 | rs144621452 | T | B | N | N | N | T | N | N | **0** |
| PCID2 | rs140418080 | T | B | - | N | - | - | N | N | **0** |
| CHD6 | rs140007009 | T | P | **D** | **D** | **M** | **D** | **D** | **D** | **6** |
| NSMCE3 | rs145058546 | **D** | B | U | **D** | **M** | T | N | **D** | **3** |
| FAM193A | rs34435530 | **D** | B | **D** | **D** | L | T | N | **D** | **4** |

### Co-segregation of *TMEM175* p.R377H with SQTS in the family

After the identification of the p.R377H mutation in *TMEM175* as the most likely causal variant, we identified a total of 7 subjects as mutation carriers (6 tested and 1 obligate carrier) with a high likelihood of SQTS. There were 2 additional likely carriers (I.2 and III.10), but they could not be tested as their DNA samples were not available (**Table 1**). All carriers except one exhibited a short QT interval on ECG (321±43 ms, n = 8 including all carriers and likely carriers with available data; 321.63±46 ms in carriers only, n=7 with available data), while none of the non-carriers (n=8) displayed an abnormally short QT (**Table 1 and Supplemental Table 2**). One mutation-positive individual was unaffected, indicating incomplete penetrance. Nevertheless, phenotype segregation was consistent with an autosomal dominant inheritance. Co-segregation was assessed using singlepoint parametric LOD score analysis (θ=0) with the *paramlink2* R package, assuming an autosomal dominant model with disease-allele frequency of 1×10^-4^ and a priori fixed penetrance of 0.90, consistent with incomplete penetrance for autosomal dominant channelopathies. This analysis yielded a LOD score of 1.78, below the threshold for definitive linkage (LOD>3), but provides substantial supportive genetic evidence for this extremely rare and lethal syndrome.

### Population level evidence for the role of *TMEM175* in QT interval duration

We further strengthened the genetic evidence by including a large population burden testing of *TMEM175* in QTc interval duration in the UK Biobank, which integrates WGS data and linked clinical records from approximately 500,000 individuals. We identified four carriers of the rs751828848 (chr4:958111:G>A) variant in the UK Biobank (4/500,000), however, ECG data were unavailable for these individuals and UK Biobank patient-level data protection policies prevent us from sharing specific phenotype data from carriers. Given the limited power of single-variant analyses for ultra-rare variants and the absence of ECG data for these carriers, we performed gene-based burden testing to assess *TMEM175*’s association with QT interval modulation. Among ∼118,000 UK Biobank participants with available ECG data, we evaluated 54 combinations of functional masks^14^ and allele–frequency thresholds. Rare-variant burden analyses revealed a significant association between *TMEM175* and the inverse rank-normalized QTc interval (Cauchy_ALL p = 0.0103; Cauchy_MISSENSE p = 0.00434). The strongest individual signal came from the canonical missense (AlphaMissense_PP) mask at AAF ≤1×10^-5^ (44 carriers; p = 8.80×10^-4^; β =−0.467; SE = 0.140) which would be significant even after multiple-testing correction for the 54 masks included in the analysis (p<0.05/54=9.2×10^-4^ cut-off). Sensitivity analyses using raw (non-normalised) QTc values across the same 54 combinations produced highly-similar results (Cauchy_ALL p = 0.0107; Cauchy_MISSENSE p = 0.00479), with the same leading mask (AlphaMissense_PP; n = 44; p = 1.03×10^-3^; β = −11.10 ms; SE = 3.38). Thus, rare *TMEM175* missense variation is significantly associated with an ∼11 ms (or ∼0.5 SD) shortening of the QTc interval in the general population, in agreement with the R377H variant identified by WES in this family. Notably, an 11ms reduction in QTc constitutes a substantial magnitude effect, comparable with the lower end of known LOF variants for long-QT syndrome (ranging from +10ms^15^ to +30ms^16^).

### Short QT intervals in transgenic TMEM175^R377H^ mice

To confirm the association of the TMEM175^R377H^ variant with the patientś SQTS diagnosis we developed a transgenic TMEM175^R377H^ mouse model using CRISPR-Cas9 technology (see *Supplementary Methods* for details). Similar to the SQTS patients, histology (H&E, Masson trichrome) (**Supplemental Figure 3A**), heart weight-to-body weight (HW/BW) ratio determination (**Supplemental Figure 3B**) and echocardiography showed no differences in cardiac morphology/structure and contractile function in TMEM175^R377H^ versus TMEM175^WT^ (**Supplemental Figure 3C**). On surface ECG, both male and female TMEM175^R377H^ mice exhibited clear, sex-independent short QT and QTc intervals, with no significant changes in intra-atrial (P-wave), atrioventricular (PR interval) and intraventricular (QRS) conduction (**Figure 2A and 2B**). Additionally, TMEM175^R377H^ mice exhibited a shallow QT/RR slope compared to TMEM175^WT^, reflecting a reduced QT interval adaptation to changes in heart rate (**Supplemental Figure 4**). Treatment with quinidine (Q), which has been shown to be effective in preventing life-threatening arrhythmic events in patients with SQTS^17^, failed to restore the QT and QTc intervals in the III.9 proband, while in TMEM175^R377H^ male mice restored QT levels (**Supplemental Figure 5**). This discrepancy may reflect species-specific differences in cardiac electrophysiology and drug response, as well as underlying mechanistic heterogeneity of short QT syndrome in humans compared with murine models.

**Figure 2.**
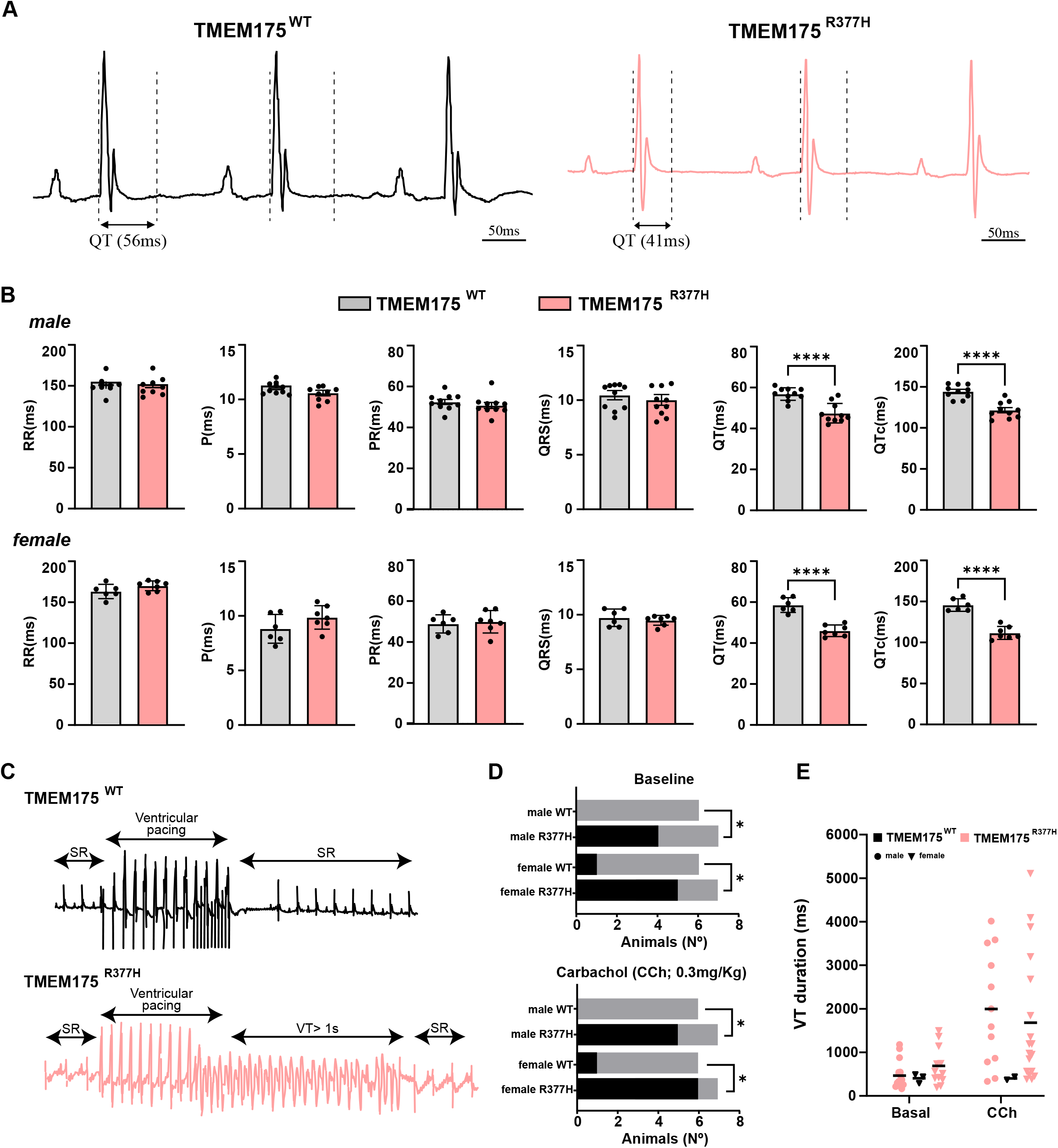
Short QT interval and increased arrhythmia inducibility in male and female TMEM175^R377H^ mice. **A)** Representative lead-II ECGs from TMEM175^WT^ (l*eft*; black) and TMEM175^R377H^ (*right*; red) mice. Note the extreme sex-independent QT interval abbreviation in TMEM175^R377H^ mice. **B)** Quantification of RR, P, PR, QRS, QT and QTc (corrected by Bazett’s formula) durations in male and female TMEM175^WT^ and TMEM175^R377H^ mice. QT and QTc interval were reduced in TMEM175^R377H^ compared to control mice (N=10 animals per group; ten consecutive beats per animal; unpaired t-test, \**P<*0.05, \*\*\*\**P<*0.0001). **C)** Representative ECG lead-II traces after a train of intracardiac ventricular pulses in TMEM175^WT^ (top; N=6) and TMEM175^R377H^ (bottom; N=7). Note the induction of long-lasting VT (longer than a second) in the latter. **D)** Contingency plots of the number of animals with arrhythmogenic response after intracardiac stimulation at baseline (top) and under carbachol (CCh; 0.3 mg/Kg) (bottom). E) VT duration in TMEM175^WT^ (black) and TMEM175^R377H^ (red) mice, both at baseline (left) and under CCh administration (right). Each value is the mean ± SEM; statistical analysis using Chi-square test, \**P<*0.05.

**Figure 3.**
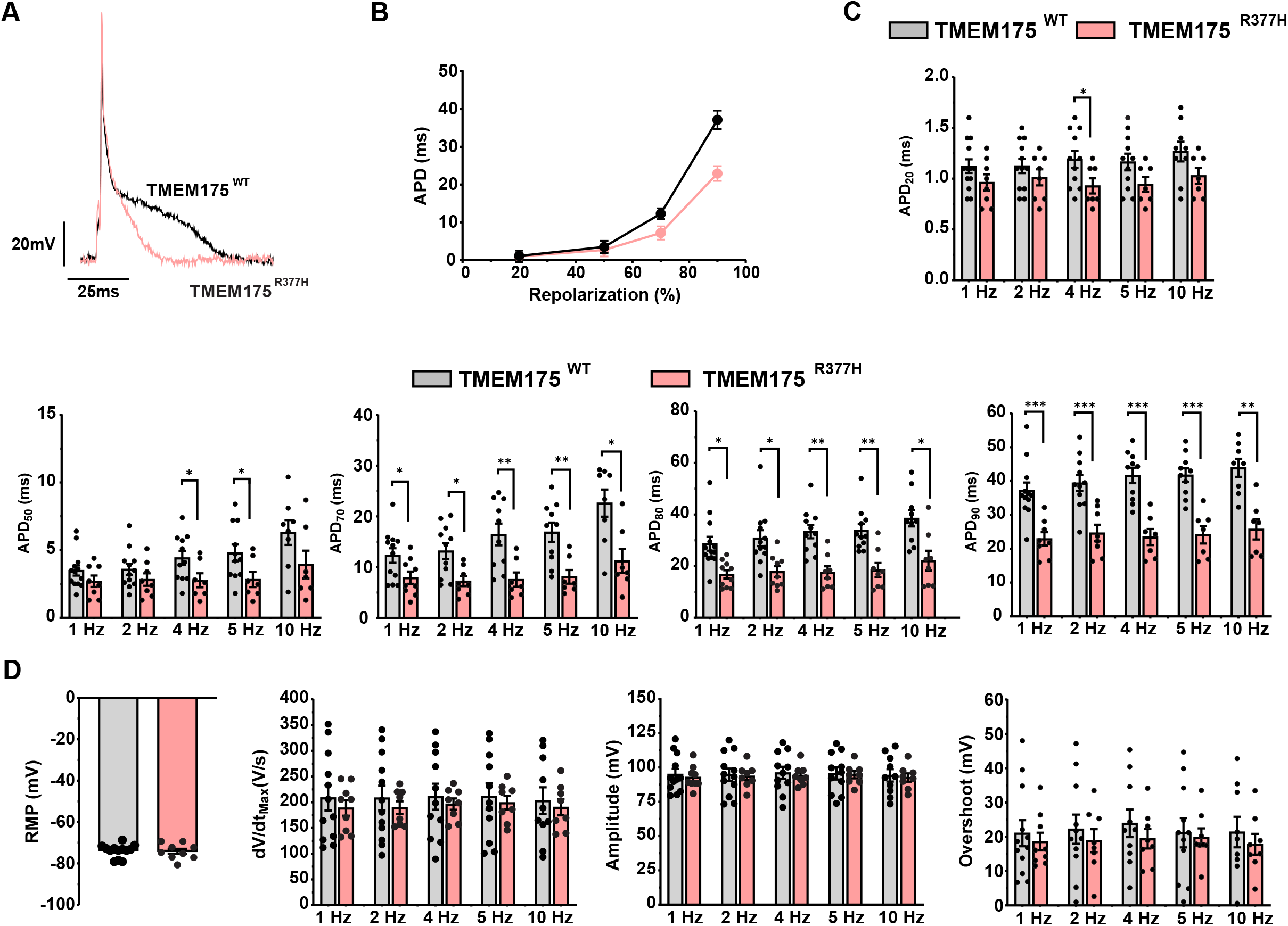
Action potential duration is abbreviated in TMEM175^R377H^ mouse cardiomyocytes. **A)** Superimposed action potentials from TMEM175^WT^ (black) and TMEM175^R377H^ (red) cardiomyocytes paced at 1 Hz. **B)** Mean APDs at 20, 50, 70 and 90% repolarization at 4 Hz pacing. APD shortening in TMEM175^R377H^ occurs from APD_50_ onwards and reaches 50% at APD_90_ (41.65±2.293 ms *vs* 23.46±2.381 ms in TMEM175^WT^ and TMEM175^R377H^, respectively). **C)** Comparison of APD in TMEM175^WT^ and TMEM175^R377H^ at 20, 50, 70, 80 and 90% repolarization at 1, 2, 4, 5 and 10 Hz (N=6 animals per group). **D)** Comparison of resting membrane potential (RMP), *d*V/*d*T, amplitude and overshoot (N=3, n=913); Each value is the mean ± SEM. Unpaired two-tailed student’s t-test or MannWhitney test were applied (\*\*\**P*<0.001, \*\**P*<0.01, \**P<*0.05).

**Figure 4.**
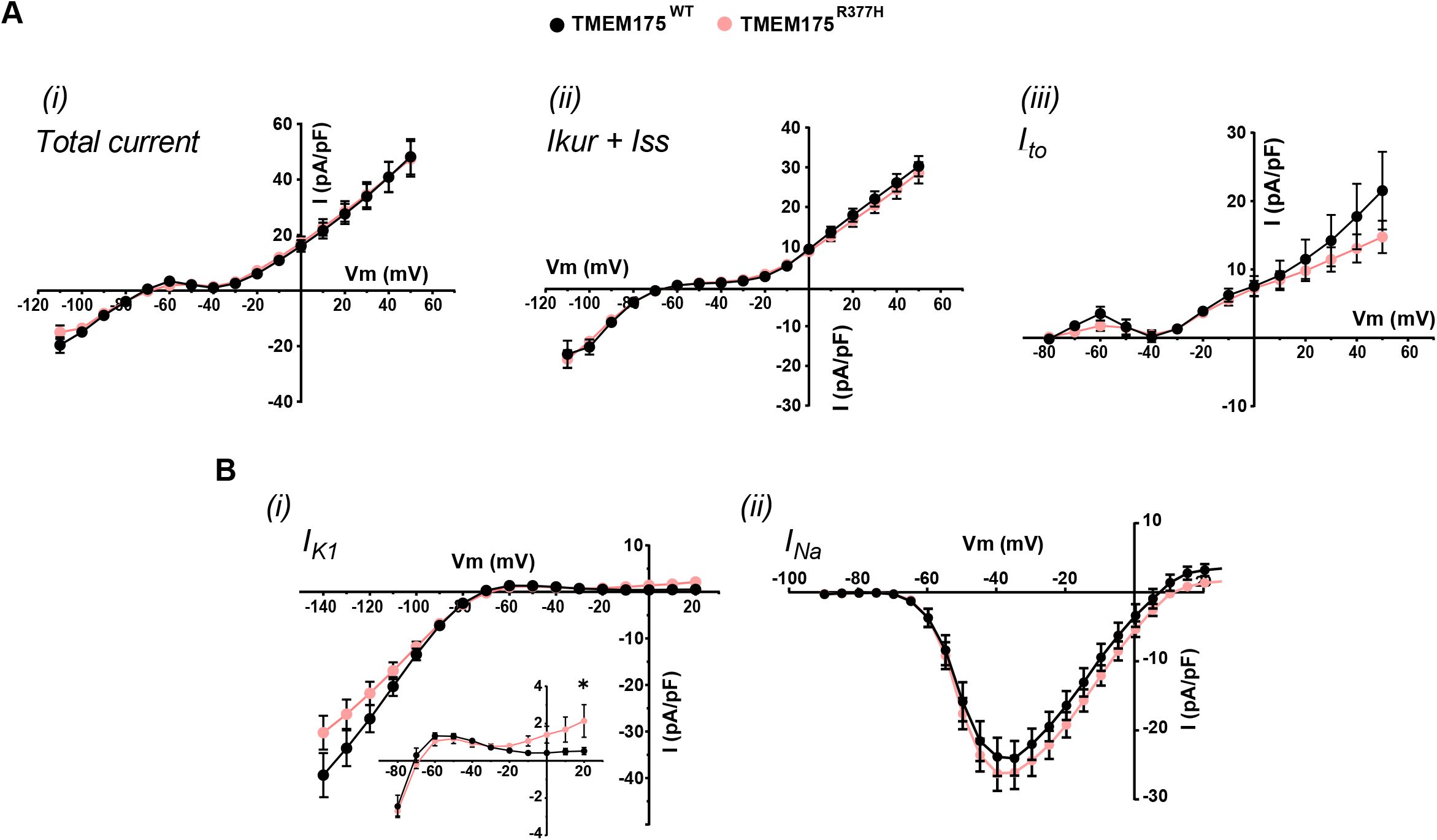
K^+^ and Na^+^ currents in TMEM175^WT^ vs TMEM175^R377H^ cardiomyocytes. A) Total I_K_ (i), I_ku_r+I_ss_ (ii), and I_to_ (iii) current-voltage (I/V) plots. **B)** TMEM175^R377H^ outward I_K1_ is significantly increased at +20 mV (i). I_Na_ I/V is unchanged (ii). (N= 3, n=9-14. Each value is the mean ±SEM; two-way ANOVA; \**P<*0.05).

### APD is abbreviated throughout the TMEM175^R377H^ mouse heart

We conducted optical mapping experiments to compare the spatial APD distribution and its dispersion throughout the ventricular epicardium of Langendorff-perfused **TMEM175^R377H^** versus **TMEM175^WT^** male mouse hearts. **Supplemental Figure 6A** shows single pixel recordings (upper) and high-resolution APD maps (lower) obtained at pacing basic cycle lengths (BCLs) of 150-250 ms. In these hearts, the largest differences in APD occurred at 50 and 80% repolarization, with substantially more abbreviation near the right ventricular base, a dispersion that may contribute to arrhythmogenesis at high activation rates (**Supplemental Figure 6B-C**). Conduction velocity (CV) maps and CV restitution curves presented in **Supplemental Figure 6D** demonstrate that the mutation did not affect CV at any BCL.

### Arrhythmias are highly inducible in TMEM175^R377H^ hearts

Despite the absence of obvious structural or functional abnormalities, TMEM175^R377H^ mice exhibited a reduced survival rate compared to TMEM175^WT^ mice. Additionally, TMEM175^R377H^ mice had fewer births and lower pup weaning rates than their TMEM175^WT^ counterparts. (**Supplemental Figure 7A-B**). While we cannot guarantee such differences were cardiac-related, evaluation of embryonic lethality in deceased TMEM175^R377H^ mice demonstrated normal development of the body organs, particularly the nervous system (**Supplemental Figure 7C**), which reinforces the concept of cardiac-related (arrhythmic) death, as previously described^18^. In addition, intracardiac ventricular stimulation using consecutive trains of stimuli at 10 and 25 Hz showed a higher arrhythmia susceptibility in both male and female adult TMEM175^R377H^ mice. Under baseline conditions, 4 out of 7 (57%) male and 5 out of 7 (71%) female TMEM175^R377H^ mice developed VT/VF (**Figure 2C and 2D**), some lasting >1 s (**Figure 2E**), compared with 0 out of 6 (0%) in both control male and female TMEM175^WT^ mice. Based on the proband’s family history of SCD while resting, we tested arrhythmia inducibility in mice under parasympathetic stimulation with carbachol (0.3 mg/Kg). Notably, male and female TMEM175^R377H^ mice exhibited increased arrhythmia inducibility and VT duration (5 out of 7 male, 71%; 6 out of 7 female, 85%) compared with control mice (0 out of 6 male, 0%; 1 out of 6 female, 16%) (**Figure 2D and 2E**), supporting a role of vagal influence in the arrhythmogenic phenotype. Altogether, the results indicate that the TMEM175^R377H^ mutation recapitulates the SQTS phenotype of the family, establishing a sex-independent arrhythmogenic substrate and a new form of SQTS. These data are compatible with the SCD in 2 carrier relatives of the proband (II.5 and III.10; **Figure 1**).

### APD is abbreviated in isolated TMEM175^R377H^ cardiomyocytes

Current-clamp recordings in isolated TMEM175^R377H^ cardiomyocytes revealed consistent and significantly abbreviated APDs at all frequencies tested, particularly during the final phase of AP repolarization (**Figure 3A-C**). The mean APD_90_ in TMEM175^R377H^ was about 50% shorter than TMEM175^WT^ compatible with the short QT phenotype of the patients and TMEM175^R377H^ mice (**Figure 3 C and Supplemental Table 3**). As shown in **Figure 3D**, RMP, *d*V/*d*t, AP amplitude and overshoot were not affected by the mutation.

In voltage clamp experiments, we carefully determined potassium and sodium current densities in freshly isolated TMEM175^WT^ and TMEM175^R377H^ cardiomyocytes. **Figure 4A** and **Supplemental Figure 8** show superimposed IV relationships and current traces, respectively, for total K^+^ current as well as I_kur_+I_ss_, and I_to_. There were no remarkable differences between the two genotypes. In **Figure 4B(i)** the outward component of the barium-sensitive inward rectifier potassium current (I_K1_) of TMEM175^R377H^ cardiomyocytes was slightly but significantly larger at +20 mV. On the other hand, neither the sodium current (I_Na_) nor its activation and inactivation voltage-dependence exhibited any significant changes (**Figure 4B(ii) and Supplemental Figure 9).** In summary, among all the outward and inward currents surveyed, only the outward component of I_K1_ showed an increase. However, we were unconvinced this small repolarizing current alone was sufficient to explain the APD and QTc shortening phenotype.

### SQTS patient-specific iPSC-CMs monolayers recapitulate the shortened APD phenotype

From the foregoing, the transgenic TMEM175^R377H^ mouse model recapitulates the patient’s SQTS phenotype (**Table 1**). We next used patient-specific iPSC-CMs to validate our animal results in a human cell model, considered as a “clinical trial in a dish”^19^. We first established the patient-specific R377H cell line and then generated an isogenic cell line by CRISPR-Cas9 (see Supplemental *Methods*). The isogenic line acted as the most accurate control with the patientspecific genetic background, establishing a causal relationship of the mutation. Pluripotency of patient-specific and corrected iPSCs was confirmed before starting the experiments (**Supplemental Figure 10**). We performed optical mapping experiments in monolayers paced at BCLs between 600 and 1000 ms. APD was shorter in the patient-specific R377H cell line. APD_80_ at BCL 1000 ms was particularly remarkable, but there were also significant changes at APD_50_ at most frequencies tested **(Figure 5A-C and Supplemental Table 4**). As shown in **Figure 5D**, activation times and CVs were similar in patient-specific R377H and isogenic control cell monolayers. However, the CV restitution curve of the mutant monolayers was flatter than the isogenic monolayers indicative that the former adapted poorly to changes in frequency, which is a sign of pro-arrhythmia typical in SQTS cases (**Figure 5D, right**). Importantly, TMEM175^R377H^ monolayers exhibited a higher incidence of rotor activity compared to isogenic controls (**Figure 5E-F**). In addition, several mutant monolayers displayed spatial dispersion of refractoriness, with intermittent 2:1 conduction block (**Figure 5G and Supplemental video 1-3**). These conduction abnormalities increased heterogeneity in wave propagation, potentially promoting re-entry and arrhythmogenic risk.

**Figure 5.**
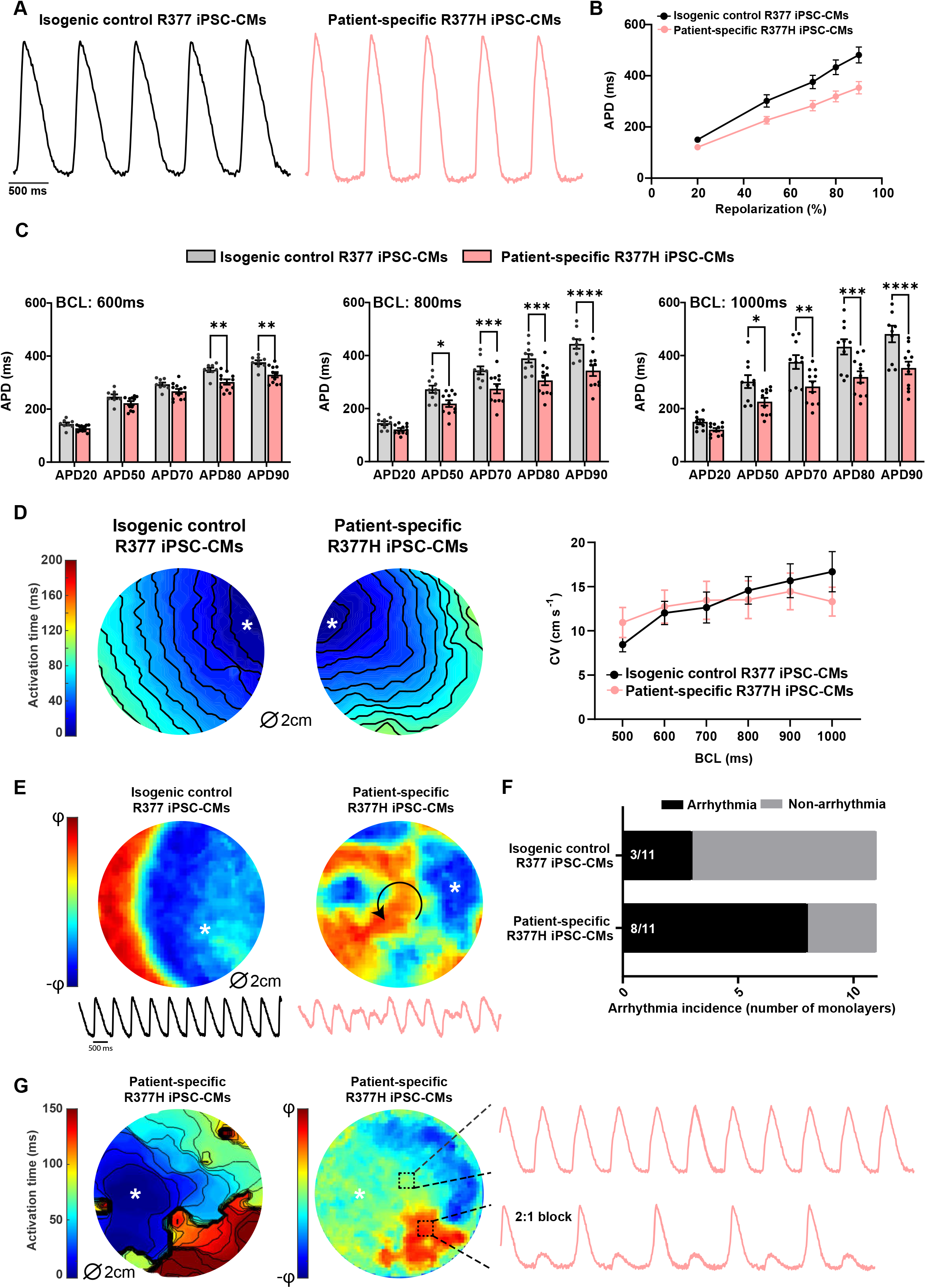
APD is abbreviated in patient-specific R377H iPSC-CM monolayers. **A)** Representative optical AP traces at BCL 800 ms from corrected R377 isogenic control (top left) and patient-specific R377H iPSC-CM (top right) monolayers. **B)** Mean APDs at 20, 50, 70, 80 and 90% repolarization at BCL 800 ms pacing. **C)** Bar graphs compare the mean APDs at 20, 50, 70, 80 and 90% repolarization at BCLs 600, 800, and 1000 ms. Note the significant APD_80_ abbreviation in R377H iPSC-CM monolayers at all frequencies. **D)** Left, Activation maps with 10 ms-isochrones from isogenic control (left) and patient-specific R377H iPSC-CM monolayers (right) paced at a BCL of 1000 ms. White asterisks indicate pacing sites. Right, Superimposed CV restitution curves. **E)** Phase maps and single-pixel recordings show normal wave propagation in representative isogenic iPSC-CM and the initiation of a re-entrant arrhythmia in patient-specific R377H iPSC-CM monolayers. The white asterisk indicates the site of impulse initiation. The color bar indicates the AP phase. **F)** Contingency plots show greater arrhythmia inducibility in mutant vs control iPSC-CM monolayers. Each value is the mean ± SEM (Fisher’s exact test for contingency data). **G)** Representative isochronal (left) and phase (middle) maps of an R377H iPSC-CM monolayer showing an area of intermittent 2:1 block. On the left are single-pixel recordings from the same monolayer illustrating the local 2:1 block pattern (bottom). White asterisks indicate the site of impulse initiation (N=5 n=11; 2-way ANOVA; \*\**P*<0.01). Each value is the mean ±SEM.

Altogether, both human and murine models fully recapitulated the patientś phenotype, providing strong evidence for an association between the *TMEM175* gene and SQTS. These findings suggest that altered frequency-dependent conduction dynamics in TMEM175^R377H^ monolayers may act as a critical trigger for SCD in this family.

### R377H does not disturb TMEM175 protein expression, structure or stability

Total protein analysis demonstrated that isolated mouse cardiomyocytes expressing TMEM175^WT^ and TMEM175^R377H^ have similar TMEM175 protein expression levels (**Supplemental Figure 11A**). In addition, quantitative real-time PCR (qRT-PCR) showed a similar mRNA expression level of the *TMEM175* gene (**Supplemental Figure 11B**), indicating that our models did not operate in haploinsufficiency. Importantly, immunolocalization studies further showed TMEM175 displaying a similar subcellular distribution pattern in both wild-type and mutant cardiomyocytes. Co-staining with the lysosomal marker LAMP1 demonstrated no detectable differences in co-localization (**Supplemental Figure 11C**), supporting the conclusion that the R377H variant does not impair lysosomal targeting of TMEM175. Arg377 localizes at the 4^th^ transmembrane domain of the second repeat (**Supplemental Figure 11D**). Alignment of TMEM175 sequences in the Arg 377 region from various species shows that residue 377 is highly conserved, Arginine (R) being the amino acidic residue expressed in all analyzed mammals, including humans and mice (**Supplemental Figure 11E**). Thus, we used *in-silico* homology modeling to derive predictions of the molecular structure of the TMEM175^R377H^ mutant channel to understand the possible mechanisms underlying its dysfunction. In **Supplemental Figure 11F**, no remarkable changes are observed in either the pore or the overall architecture of the channel. Atomic level modeling showed that, compared to the WT channel, TMEM175^R377H^ undergoes a slight reorganization in its structure (TMscore 0.91; RDMS: ∼2,21Å for homodimer or TMscore 0.95; RDMS: ∼2,08Å for heterodimer, **Supplemental Figure 11G**). Similarly, the Gibbs free-energy values for TMEM175^R377H^ were similar to WT (TMEM175^WT^: -2772,376 vs TMEM175^R377H^: -2899,055 and 2890,050 for homo and heterodimer respectively) (**Supplemental Figure 11G**). Therefore, although the mutation is located at an evolutionary conserved residue, it does not affect the overall structure of the channel or its stability and expression levels. This paradoxical result led us to study already known TMEM175 channel functions to elucidate the TMEM175^R377H^ effect.

### Mitochondria are disorganized and dysfunctional in TMEM175^R377H^ cardiomyocytes

Further lysosomal analysis revealed an increased expression of the lysosomal marker Lamp1 and elevated LC3-II/LC3-I ratio in isolated TMEM175^R377H^ cardiomyocytes compared to control (**Supplemental Figure 12A**). We also observed an increase in total LC3 expression (**Supplemental Figure 12B**), which strongly suggested an alteration of the normal autophagy dynamics. To directly assess the level of autophagic flux, we expressed the tandem fluorescent reporter mRFP-GFP-LC3 via AAV in cardiomyocytes. Mutant cells exhibited a reduced GFP/RFP ratio, consistent with GFP quenching in autolysosomes, indicating that autophagosomes efficiently fuse with lysosomes and that autophagic flux remains active compared to control cells (**Supplemental Figure 12C**). We also evaluated mitophagic activity by analyzing Parkin protein levels, given that Parkin deficiency has been shown to enhance autophagy and mitophagy induction^20^. In TMEM175^R377H^ cardiomyocytes, Parkin expression was similar to controls (**Supplemental Figure 12D**), suggesting that Parkin-mediated mitophagy remains intact. Together, these findings support the role of TMEM175 in maintaining mitochondrial quality control. However, molecular mechanisms were independent of changes in intracellular pH, as evidenced by fluorescence microscopy experiments (**Supplemental Figure 12E**).

Ultrastructural analysis using transmission electron microscopy (TEM) showed similar lysosomal number and size in mutant cardiomyocytes (**Figure 6A-B**), which could indicate an altered lysosomal function rather than a change in structural quantity. In contrast, TEM images showed a clear disorganization of mitochondrial architecture in mutant cells **(Figures 6C-D**). Consistently, the mitochondrial marker Tom20 was highly expressed in TMEM175^R377H^ cardiomyocytes (**Figure 6E**), and qRT-PCR experiments showed a non-significant tendency for an increase in mitochondrial genome copy number (**Figure 6F**). To further characterize mitochondrial function and homeostasis, we assessed mitochondrial membrane potential (ΔΨm) and mitochondrial superoxide production using MitoSOX. In these studies, HEK293T cells were either cotransfected with TMEM175^WT^ or heterozygous WT/R377H construct at a 1:1 ratio, and cells were harvested 48 hours post-transfection. Flow cytometry analyses revealed elevated ROS production (**Figure 6G**) and reduced ΔΨm (**Figure 6H**) in the heterozygous conditions TMEM175^WT/R377H^. We also evaluated mitochondrial dynamics by analyzing the expression of the key fission and fusion regulators Drp1 and OPA1, respectively. Notably, adult cardiomyocytes maintained comparable expression levels of mitochondrial dynamics proteins (**Figure 6I**). Altogether, these findings indicate that the TMEM175 mutation impairs mitochondrial integrity and function, leading to disrupted cellular energy homeostasis^9^ without overt alterations in mitochondrial dynamics protein expression.

Therefore, to evaluate a potential alteration in mitochondrial function and energy homeostasis, we performed Seahorse experiments in isolated murine cardiomyocytes. The analysis revealed significant reductions in maximal respiration, spare respiratory capacity and ATP synthesis in TMEM175^R377H^ cardiomyocytes (**Figure 7A**), indicating mitochondrial dysfunction. Lysosome–mitochondria contact sites are increasingly recognized as critical hubs for interorganelle communication^21^. Although TMEM175 is not yet directly linked to the structural formation of these contact sites, it plays a well-established role in lysosomal ion homeostasis, which can influence signaling pathways that affect mitochondrial performance. It is plausible that TMEM175 activity influences mitochondrial membrane potential (ΔΨm), which is essential for ATP production via oxidative phosphorylation and is tightly linked to proton gradients across the mitochondrial inner membrane. To further explore this potential connection, we assessed mitochondrial membrane potential and found that TMEM175^R377H^-expressing HEK293T cells exhibited significant depolarization (**Figure 6H**). This finding suggests that altered lysosomal ion handling may compromise mitochondrial function, possibly through dysregulated proton or potassium flux. These results further support a functional link between TMEM175^R377H^ and the lysosome–mitochondria axis, prompting us to investigate mitochondrial function. Further analysis revealed reduced ATP levels (**Figure 7B**), indicating energetic imbalance in TMEM175^R377H^ cardiomyocytes.

**Figure 6.**
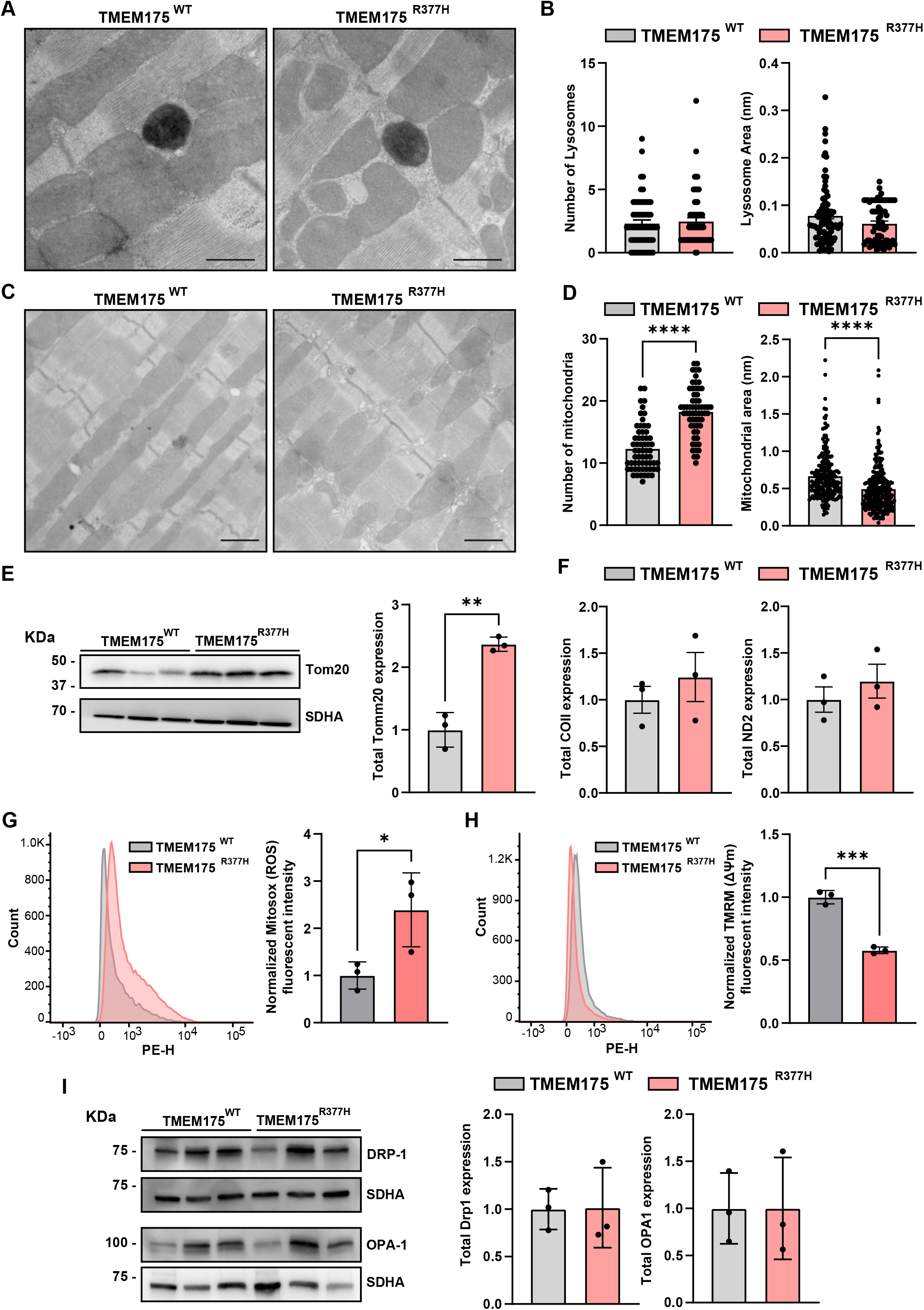
Mitochondria are abnormal in TMEM175^R377H^ cardiomyocytes. **A)** Representative Transmission electron microscopy (TEM) images from TMEM175^WT^ and TMEM175^R377H^ cardiomyocytes showing detailed lysosomes and mitochondria organelles (scale bar 10 µm). **B)** Graphs show no differences in total lysosome number (left) and lysosome area (right) in TMEM175^WT^ vs TMEM175^R377H^ cardiomyocytes (N=3 animals per group, n=15-20; Unpaired ttest). **C)** Zoom out of TEM images showing clear defects in the disposition and morphology of mitochondria in TMEM175^R377H^ cardiomyocytes. **D)** Bar plot of mitochondria number (left) and mitochondria area (right) show changes in TMEM175^R377H^ cardiomyocytes (N=3 animals per group, n=15-20; Unpaired ttest; \*\*\*\**P*<0.0001). **E)** Western blot analysis of total TOM20 protein expression in WT and mutant cardiomyocytes. Shown are TOM20 (upper band), SDHA (middle band) and the stain-free (*bottom*) protein (*left)*, along with the quantification of TOM20 protein levels relative to SDHA (*right)* (N=3 animals per group; Unpaired t-test \*\**P*<0.01). **F)** qRT-PCR quantification of total mitochondrial cytochrome oxidase II (COII; *left*) and NADH dehydrogenase 2 (ND2; *right*) mRNA expression. DNA mitochondrial to nuclear genome copies ratio (N=3; mean ± SEM; Unpaired t-test). **G)** Quantification of mitochondrial superoxide production by MitoSOX staining (5µM) in HEK293T cells transfected with TMEM175^WT^ and TMEM175^WT/R377H^ expressing plasmid by flow cytometry. Representative histograms show the distribution of MitoSOX fluorescence and bar graphs represent the mean fluorescence intensity normalized control levels (N=3; mean ± SEM; Unpaired t-test; * P<0.05). **H)** Assessment of mitochondrial membrane potential (ΔѰm) using tetramethylrhodamine methyl ester (TMRM, 10nM) in HEK293T cells transfected with a TMEM175^WT^ and TMEM175 ^WT/R377H^ expressing plasmid by flow cytometry. Representative histograms illustrate the distribution of TMRM fluorescence and quantification of mean fluorescence intensity is shown as relative values normalized to control (N=3; mean ± SEM; Unpaired t-test; \*\**P*<0.001). **I)** Evaluation of mitochondrial dynamics in TMEM175^WT^ and TMEM175^R377H^ mouse cardiomyocytes. Protein expression levels of the mitochondrial fission marker dynamin-related protein 1 (Drp1) and the fusion marker optic atrophy protein 1 (Opa1) were analyzed by Western blot and normalized to succinate dehydrogenase complex flavoprotein subunit (SDHA). Quantification of protein expression is presented as bar graphs showing relative levels compared to TMEM175^WT^ (N=3; mean ± SEM; Unpaired t-test).

**Figure 7.**
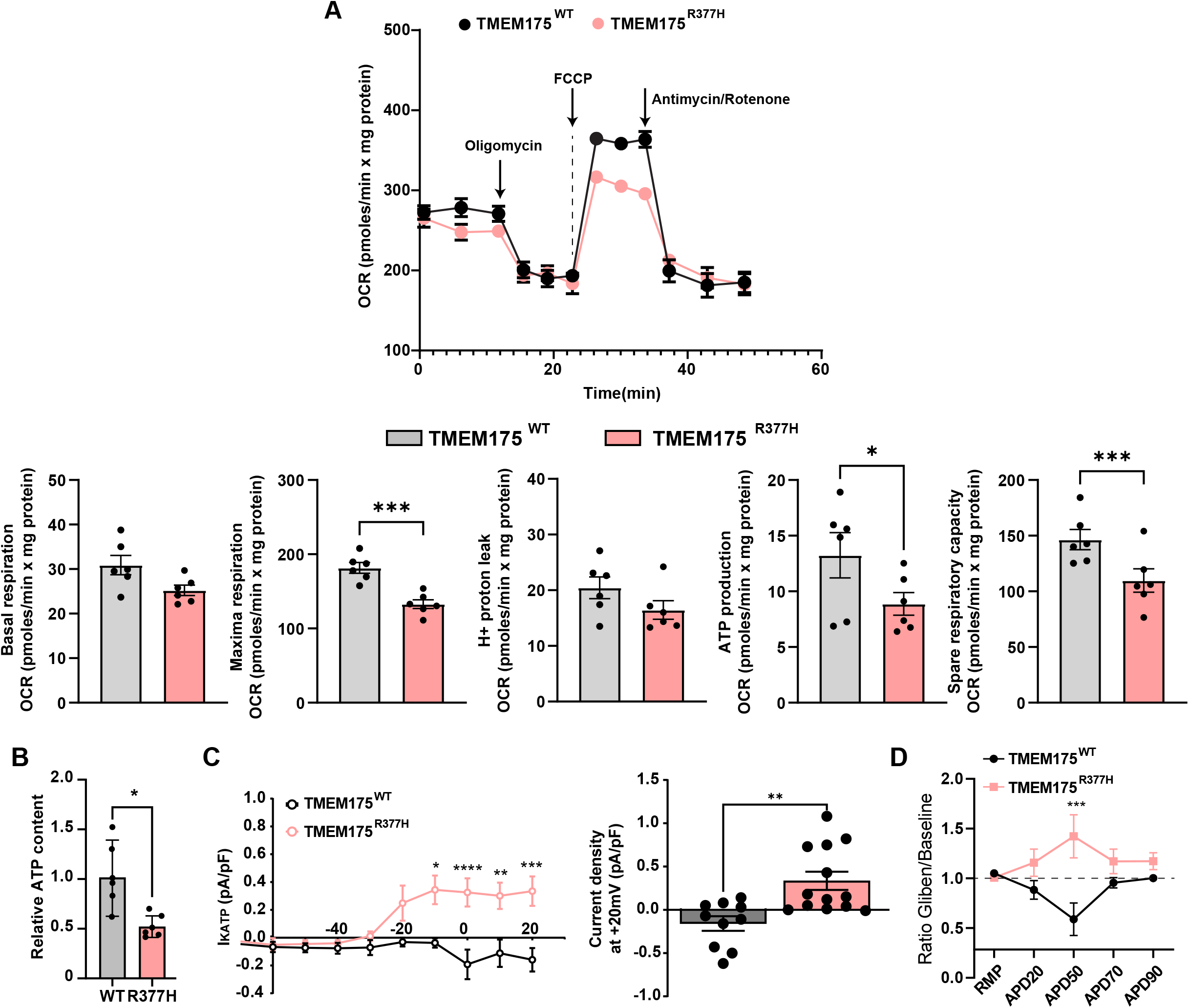
TMEM175^R377H^ cardiomyocytes have reduced mitochondrial ATP production coupled with I_KATP_ activation. **A)** TMEM175^R377H^ showed compromised mitochondrial respiration capacity and decreased maximal mitochondrial OCR and ATP levels (N=5; 2-way ANOVA; \**P<*0.05, \*\*\**P*<0.001). **B)** Quantifying the relative ATP content measured with fluorometry in TMEM175^WT^ and TMEM175^R377H^ isolated cardiomyocytes (N=6; Unpaired t-test; \**P<*0.05). **C)** I/V relationship of the ATP-sensitive current (I_KATP_) recorded in the perforated patch configuration shows a significant activation of I_KATP_ at more positive voltages. Graph shows current density at +20 mV (mean ±SEM; N=3, n=10-13; 2-way ANOVA; \**P<*0.05, \*\**P*<0.01, \*\*\**P*<0.001, \*\*\*\**P<*0.0001). **D)** Action potential recordings under I_KATP_-inhibitor glibenclamide (1 µM). The graph shows the ratio of APD in Glibenclamide and Baseline.

### Low ATP activates I_KATP_ in TMEM175^R377H^ mice

One of the major changes produced by metabolic impairment of ventricular cardiomyocytes is APD abbreviation, reducing the duration of the refractory period and increasing the risk of re-entrant arrhythmias^22,23^. Thus, the low ATP levels and the absence of major modifications in the canonical potassium and sodium currents that could justify the short QT phenotype led us to study the ATPsensitive current (I_KATP_), which is inactivated by ATP and has already been associated with APD shortening^24,25^. Patch-clamp experiments revealed an increased I_KATP_ in mutant cardiomyocytes, starting from -20 mV and reaching its maximum at +20 mV. In contrast, control cardiomyocytes lacked measurable I_KATP_ (**Figure 7C**). To further demonstrate that I_KATP_ is the principal mediator of APD shortening, we measured APD in the presence of glibenclamide, a well-known I_KATP_ inhibitor. Glibenclamide infusion (1 µM) had no effect in isolated TMEM175^WT^ cardiomyocytes but clearly increased APD in TMEM175^R377H^ cardiomyocytes, confirming the key role of I_KATP_ (**Figure 7D**). The results indicate that the energy imbalance and ATP reduction produced by TMEM175^R377H^-induced disruption of the lysosomal-mitochondria axis activates sarcolemmal I_KATP_ and shortens the APD.

### TMEM175^R377H^ disrupts Bcl-2-mediated lysosomal-mitochondrial interaction

There is no consensus on how the TMEM175 channel affects mitochondrial activity^9^. However, physical lysosome-mitochondrial interaction has been extensively described in cardiomyocytes and is presumed to be a crucial factor in maintaining the energetic state of the cell^26–29^. Interestingly, TEM images revealed a significant reduction in lysosome-mitochondria interaction in TMEM175^R377H^ murine cardiomyocytes. This was evidenced by an increased average lysosomemitochondria distance and a decreased contact perimeter between both organelles **(Figure 8A-B**). These findings indicate that the R377H mutation disrupts the lysosome-mitochondrial regulatory axis.

**Figure 8.**
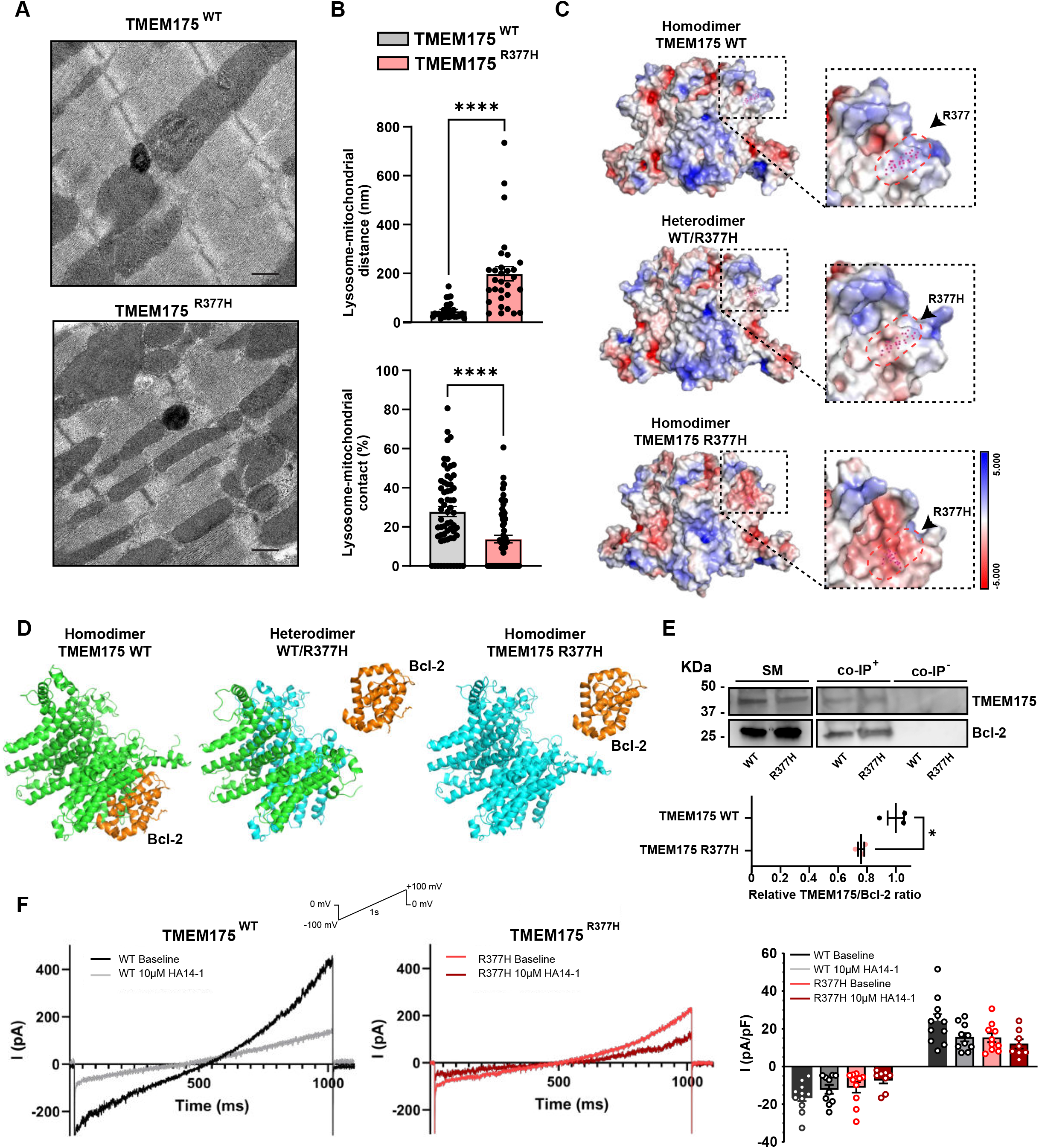
TMEM175^R377H^ disrupts the lysosome-mitochondria regulatory axis. **A)** Representative TEM images show impairment of lysosome-mitochondria interaction in TMEM175^R377H^ cardiomyocytes (scale bar 5 µm). **B)** Quantification of the lysosome-mitochondria distance (nm) and the lysosome-mitochondria perimeter of contact in both conditions (mean ±SEM; N=3, n=15-20; Unpaired t-test; ***P<0.0001). **C)** Representative in-silico electrostatic surface maps of TMEM175^WT^, TMEM175^WT/R377H^ heterodimer and TMEM175^R377H^ homodimer channels. A detailed electrostatic change is shown for the 377-residue surrounding area in each condition (black arrowheads). The color bar indicates an energetic transition from positive (blue) to negative charge (red). **D)** In-silico ligand-docking analysis between Bcl-2 protein and TMEM175^WT^, TMEM175^WT /R377H^ heterodimer and TMEM175^R377H^ homodimer channels. Note that heteroand homozygous conditions completely lack Bcl-2 binding capacity. **E)** Immunoprecipitation of Bcl-2 protein with TMEM175^WT^ or TMEM175^R377H^ channel in HEK293T expressing cells. IP, immunoprecipitating antibody; WB, immunoblotting antibody (N=3; independent replicates). **F)** Representative whole-cell recording of total current in the absence and the presence of 10 μM of Bcl-2-selective inhibitors HA14-1 from TMEM175^WT^ and TMEM175 ^WT/R377H^ transfected HEK293T cells. The graph shows normalized peak currents (I/I0) from -100 to 100 mV for each condition (N=4 experiments, n=8 replicates per group).

To elucidate the molecular mechanism underlying this loss, we examined the electrostatic charges in the atomic environment surrounding the Arg377 residue. Specifically, the R377H mutation caused a significant shift from a positive to an extremely negative charge, potentially affecting the interaction with accessory proteins in both the TMEM175^WT/R377H^ heterodimer and TMEM175^R377H^ homodimer (**Figure 8C**). Interactions between TMEM175 and mitochondrial proteins like succinate dehydrogenase complex flavoprotein subunit A (SDHA), ATP Synthase F1 Subunit Gamma (ATP5C1)^30^ and B-cell leukemia/lymphoma 2 protein (Bcl-2) have been previously described^31^. Recent studies specifically highlight a relationship between TMEM175 and Bcl-2, interacting through the Arg377 residue^31^, where our mutation is located. To predict the viability of TMEM175-Bcl2 interactions, we conducted a molecular docking simulation. Our results revealed that TMEM175^R377H^ disrupts this interaction, following a dominant-negative pattern. Unlike the TMEM175^WT^ channel, neither the TMEM175^WT/R377H^ heterodimer nor TMEM175^R377H^ homodimer exhibited physical interaction with Bcl-2 (**Figure 8D**). To experimentally validate these findings, we conducted co-immunoprecipitation assays in HEK293T cells. Bcl-2 was co-transfected with either TMEM175^WT^ or a heterozygous WT/R377H construct at a 1:1 ratio, and cells were harvested 48 hours post-transfection for protein immunoprecipitation and SDS-PAGE analysis. Western blot results confirmed that TMEM175 physically interacts with Bcl-2. Importantly, TMEM175^R377H^ exhibited reduced binding compared to TMEM175^WT^ when normalized to immunoprecipitated Bcl-2, supporting the prediction that the mutation weakens this interaction (**Figure 8E**). Together, these data indicate that the R377H variant disrupts TMEM175–Bcl-2 interaction, which may contribute to altered lysosome–mitochondria contacts and downstream cellular dysfunction. To further confirm the loss of TMEM175-Bcl-2 interaction, we evaluated the K^+^ flux capacity of overexpressed TMEM175 channels in HEK293T in the presence of the Bcl-2 inhibitor HA14-1^31^. Patch-clamp experiments showed that, at baseline, mutant channels exhibited reduced total current compared to control. More notably, while HA14-1 treatment (10 μM) decreased TMEM175^WT^ channel current by 40%, the TMEM175^R377H^ mutant channel showed minimal reduction **(Figure 8F**). These findings strongly indicate that TMEM175^R377H^ significantly reduces Bcl-2 binding capacity, potentially disrupting lysosomal-mitochondrial interactions.

## Discussion

This study identifies the first potential SQTS-associated variant in the endolysosomal TMEM175 ion channel, and provides evidence supporting its possible involvement in the pathogenesis of SQTS and SCD. We explored a novel lysosome-mitochondria axis as a key modulator of APD. TMEM175^R377H^ disrupts this axis, impairing mitochondrial ATP production and activating I_KATP_, which shortens APD and the QT interval, increasing the risk of arrhythmias. Despite incomplete penetrance and a LOD score below the threshold for definitive linkage, familial segregation supported autosomal dominant inheritance, while population-level burden analyses in the UK Biobank provided independent evidence implicating TMEM175 in QTc interval modulation.

Unlike a targeted approach focused on conventional plasma membrane channels, our unbiased strategy in a family with high incidence of SQTS identified TMEM175, which encodes a lysosomal protein, pointing to a previously unrecognized contribution to SQTS. This finding is particularly relevant given that genetic diagnosis is currently achieved in only about 20 to 30% of patients.

The ClinGen Gene Curation Expert Panel provides a standardized approach to determining causative genes in cardiac arrhythmias, including CPVT and SQTS, assigning 12 points for genetic evidence and 6 for experimental evidence^32^. To date, only the *KCNH2* gene has a definitive assessment for SQTS, while *KCNQ1*, *KCNJ2*, and *SLC4A3* are classified as “strong” or “moderate”^32^. Other previously associated genes have been reclassified as “limited” due to insufficient supporting data. In this context, our study highlights TMEM175 p.Arg377His as a potential SQTS-associated variant. In particular, the recovery of APD in the isogenic iPSC-CM model provides the strongest functional evidence supporting its pathogenic role. By integrating familial genetic evidence, population-level burden analysis, and functional validation in *in vivo* and *in vitro* models, our study provides a comprehensive body of evidence supporting the candidacy of *TMEM175* to be evaluated by the ClinGen Expert Panel as a novel SQTS-associated gene. Nevertheless, we note that further genetic and functional studies will be required to establish a strong/definitive gene-disease relationship according to ClinGen criteria.

The discovery of TMEM175^R377H^ broadens arrhythmia genetics, demonstrating the diagnostic value of WES and extending SQTS genetics beyond traditional ion channels. Likewise, secondary SQTS has been associated with *SLC22A5* mutations^33^, which impair carnitine transport and may disrupt cardiomyocyte energy metabolism. More broadly, alterations in cellular energy homeostasis may influence cardiac electrophysiology, potentially affecting QT interval duration and arrhythmia susceptibility^34^. Our results provide solid evidence supporting an association between the lysosomal channel *TMEM175* and SQTS. Patient-specific iPSC-CMs monolayers recapitulated the shortened APD and arrhythmic phenotype. In addition, a CRISPR mouse model expressing TMEM175^R377H^ reproduced phenotypic features consistent with human SQTS clinical signs, including shortened QT and higher arrhythmia risk, in the absence of structural defects. Moreover, the mouse model recapitulated the shallow QT/RR slope observed in affected patients on Holter ECG recordings (**Supplemental Figure 1**), reflecting reduced QT interval adaptation to changes in heart rate, as previously described in classical SQTS patients with *KCNH2* mutations^12^.

TMEM175^R377H^ mice also had lower survival rates, indicating a possible link to cardiac death, though more research is needed. TMEM175^R377H^ cardiomyocytes showed APD shortening independent of Ca^2+^ dynamics. Increased I_K1_ contributed to early repolarization but I_KATP_ activation fully explained the SQTS phenotype. KATP channels, first found in heart cells^35^, carry an outward current that is usually inhibited by ATP. They are made of Kir6 and SUR subunits^36,37^. Normally closed, these channels can open via specific molecular interactions^38,39^. Our data show that the TMEM175^R377H^ mutation disrupts cell energy balance, activating I_KATP_, which is usually inactive. Once active, I_KATP_ hastens repolarization during the cardiac AP, thereby shortening the APD and causing a new type of SQTS. Increased I_K1_ also contributes. There may be additional interactions between I_KATP_ and I_K1_, but this is outside the scope of our current study^40^.

The mutation did not affect TMEM175 mRNA or protein levels. Modeling showed that TMEM175^R377H^ causes no major structural or energetic changes. However, the expression of lysosomal and autophagy markers Lamp1 and LC3 was altered, indicating lysosomal-mediated autophagy dysfunction^41^. TMEM175, once considered a K^+^ channel, now appears to mainly act as a proton (H^+^) leak channel under normal lysosomal conditions^7^. Despite lysosomal changes, intracellular pH remained stable, ruling out the SQTS mechanism seen with the AE3 exchanger^13^. These results suggest the mutation may alter lysosomal function by affecting interactions with other proteins or organelles. We also observed mitochondrial dysfunction, with increased mitochondrial mass and number. Bioenergetics assays showed reduced ATP production and total ATP content, supporting our hypothesis.

Our results suggest that mitochondrial dysfunction arises from the loss of physical interactions between lysosomal and mitochondrial proteins, disrupting the lysosome-mitochondria axis. The functional relevance of this interaction is well established. Disruption of lysosome-mitochondria coupling is associated with functional defects that significantly affect cardiac function^42–44^. Moreover, growing evidence suggests that these organellar interactions are crucial for cellular homeostasis and the stress response^29,45–47^. However, most of the molecular interconnectors involved remain undefined. In this context, TMEM175 has been shown to interact with a wide range of mitochondrial proteins, suggesting a role beyond that of a mere ion channel, potentially acting as a lysosome-mitochondria interactor^9^.

Recent studies have shown that Bcl-2 inhibits TMEM175 channel function by binding to the Arg377 residue^31^, which is important for regulating cell death and mitochondrial health in neurons. In the heart, this interaction affects organelle communication rather than cell membrane excitability. Our electron microscopy analysis revealed that the TMEM175^R377H^ mutation disrupts the lysosome-mitochondria axis in heart cells, leading to increased distance between organelles and impaired mitochondrial function, possibly due to defective autophagy. This disruption lowers ATP production and activates I_KATP_ channels, shortening the APD in cardiomyocytes^24,25^. Interestingly, our findings suggest that Bcl-2 may act as a channel activator in this context, highlighting a potentially tissue-specific regulatory effect distinct from the previously described inhibitory interaction in neurons. The TMEM175^R377H^ mutation prevents normal Bcl-2 binding, as supported by docking analysis and in vitro tests showing that a Bcl-2 inhibitor affects only the wild-type channel. It is unclear if the mutation impairs mitochondria by disrupting lysosomal signaling or through direct effects, TMEM175 may influence mitochondrial membrane potential via ion flux. Treatment with glibenclamide reversed the electrical abnormalities, supporting the idea that I_KATP_ activation due to mitochondrial dysfunction is key to QT shortening in these mutant cells. Further studies are needed to fully understand TMEM175’s role in lysosome-mitochondria communication and its impact on heart cell energy regulation.

In conclusion, this study provides the first evidence implicating TMEM175 in SQTS, identifying its role in disrupting lysosome-mitochondria communication rather than direct channel dysfunction, as a potential mechanism underlying the disease. Our findings expand the landscape of potential SQTS candidate genes, highlighting the contribution of proteins involved in the mitochondrial-lysosomal crosstalk. These results provide novel supportive evidence to consider *TMEM175* in the genetic testing for SQTS patients and emphasize the value of WES with quartet analysis in identifying new disease-associated genes, especially in a syndrome where only 20-30% of cases are genetically-positive. Therefore, we propose TMEM175 as a novel candidate gene for SQTS, representing the first putative association involving a non-sarcolemma protein.

## Materials & Methods

### Detailed descriptions are presented as Supplementary Material

#### Ethics Statements

All experimental animal procedures conformed to EU Directive 2010/63EU and Recommendation 2007/526/EC, enforced in Spanish Law under *Real Decreto 53/2013*. The local ethics committees and the Animal Protection Area of the Comunidad Autónoma de Madrid approved the animal protocols (PROEX 226.5/23 and PROEX 111.4/20). All procedures related to human samples have been approved by the Ethics Committee of Andalusia (PEIBA) and comply with the principles of the Declaration of Helsinki. All patients provided signed informed consent for the extraction of fibroblast.

#### Genetic Analysis

We combined family-based whole-exome sequencing, stringent variant prioritization, and independent UK Biobank rare-variant burden analyses in approximately 118,000 participants to provide complementary evidence supporting the involvement of TMEM175 in QTc shortening. See *Supplementary Material* for details.

#### Mice

Transgenic C57BL/6J mice were obtained by CRISPR-mediated gene editing. Mice were housed according to institutional guidelines and regulations. Mice were provided with free access to food and water. Mice aged 20-25 weeks were used in experiments.

#### Echocardiography

Transthoracic echocardiography was conducted blindly by an expert operator using a high-frequency ultrasound system (Vevo 2100, VisualSonics Inc., Canada) with a 40-MHz linear probe. Data were analyzed as described in *Supplementary Material*.

#### Surface ECG recordings

Mice were anesthetized using isoflurane inhalation (0.8-1.0% volume in oxygen) and maintained at 37°C on a heating plate. Fourlead surface ECGs were recorded for 5 minutes, using a subcutaneous 23-gauge needle electrode attached to each limb and connected to an MP36R amplifier unit (BIOPAC Systems). Data acquisition and analysis were performed blindly using AcqKnowledge 4.1 software.

#### *In-vivo* intracardiac recording and stimulation

An octopolar catheter (Science) was inserted through the jugular vein and advanced into the right ventricle as previously described^48^. Ventricular arrhythmia inducibility was assessed by applying consecutive trains at 10 Hz and 25 Hz.

#### Cardiomyocyte isolation

After euthanasia by cervical dislocation, excised hearts from TMEM175^WT^ and TMEM175^R377H^ mice were perfused in a Langendorff system and digested as described^48^ (see *Supplementary Material*).

#### Patch-clamping in isolated cardiomyocytes and HEK293T cells

Electrophysiological studies were performed in isolated cardiomyocytes from both ventricles and HEK293T cells. Whole-cell and perforated patch-clamp experiments and data analysis procedures were similar to those described previously^49–52^. External and internal solutions are described in **Supplemental Table 5.** Details are presented in the *Supplementary Material*.

#### Generation, culture, and differentiation of patient-specific iPSC-CMs

We obtained primary fibroblasts from the proband TMEM175^R377H^ carrier (III.9 in **Figure 1A**) by skin biopsy. Fibroblasts were reprogrammed to iPSC using the Sendai virus for transfection of Yamanaka’s factors: OCT4, KLF4, c-Myc, and SOX2, as previously described^53^. Patient-specific iPSCs were differentiated into iPSC-CMs using small molecules-mediated canonical Wnt pathway modulation as described^54,55^. iPSC-CMs were purified following the MACS negative selection protocol (Miltenyi Biotec Kit) and seeded for 14 days on a human cardiac matrix substrate to improve cardiomyocyte maturation. See *Supplementary Material* for details.

#### Optical mapping in isolated hearts and iPSC-CMs monolayers

Experiments were carried out as described previously^56–58^. Briefly, hearts from TMEM175^WT^ and TMEM175^R377H^ mice were isolated and connected to a Langendorff-perfusion apparatus^59^. Phase, conduction velocity (CV), activation and APD maps were constructed during pacing at basic cycle lengths (BCLs) between 150 and 250 ms. Optical mapping of iPSC-CM monolayers was conducted at BCLs between 600 and 1000 ms and CV, activation and APD maps were generated. Optical data were analyzed blindly using custom-made MATLAB software (Mathworks Inc.). See *Supplementary Material* for details.

#### Mitochondrial activity and ATP quantification

Mitochondrial activity assays and determination of mitochondrial oxygen consumption rates were carried out using seahorse experiments as previously described^60^. ATP content was measured in isolated cardiomyocytes using an ATP bioluminescence assay kit (Roche). See *Supplementary Material* for details.

## Statistical analysis

We used GraphPad Prism software versions 7.0 and 8.0. For non-Gaussian distributions, we applied the nonparametric Mann-Whitney test. We used oneor two-way ANOVA corrected by Šídák’s multiple comparisons test. We used Grubb’s test for outliers. Data are expressed as mean ± SEM, and differences are considered significant at p<0.05 (*p<0.05; **p<0.01; ***p<0.001; ****p<0.0001). Note that “N” refers to the number of mice or iPSCCM differentiations used and “n” to the number of cells analyzed per mouse or monolayers.

## ACKNOWLEDGEMENTS

We thank the CNIC Transgenesis Unit for generating the TMEM175^R377H^ transgenic mouse model. Confocal experiments were conducted at the CNIC Microscopy and Dynamic Imaging Unit. We thank the CNIC Bioinformatics Unit for generating the *in-silico* homology modelling simulations. We also thank the Pluripotent Cell Technology Unit in CNIC for their technical support in generating iPSC cell lines. We thank the CBM SO Electron Microscopy Unit. The CNIC is supported by the Instituto de Salud Carlos III (ISCIII), the Ministerio de Ciencia, Innovación y Universidades (MICIU) and the Pro CNIC Foundation, and is a Severo Ochoa Center of Excellence (grant CEX2020-001041-S funded by MICIN/AEI/10.13039/501100011033). We also gratefully acknowledge the contributions of the UK Biobanks participants, whose involvement made this research possible; this research has been conducted using the UK Biobank resource under application number 17660.

## AUTHOR CONTRIBUTION

J.M.R.R., I.M.C., J.J. and F.M.C co-designed the experiments; JMRR and IMC performed most of the experiments; J.J.J. provided clinical data; A.M. and A.I.M.M., performed cellular electrophysiology and analysis; P.S.P. and J.M.R.R. performed hiPSC-CMs experiments; J.M.R.R. and F.M. were in charge of *in-silico*

homology modelling and molecular docking studies; M.E.C. performed burden testing in UK Biobank, under supervision of S.J.J. and C.R.B.; A.I.M.M., P.S.P., P.G.S., A.T.G., E.C.B., R.M.R., F.B.J., A.M. and J.J.J., provided technical support, discussions and revisions; I.M.C, J.M.R.R, F.M.C. and J.J. co-wrote the manuscript; F.M.C. led the overall scientific direction and development of the manuscript. J.J. and A.M. provided funding, J.J. provided supervision, funding and revisions; All authors discussed the results and commented on and approved the manuscript.

## CORRESPONDING AUTHORS

Correspondence Francisco M. Cruz

## SUPPLEMENTAL INFORMATION

Extended Materials & Methods

Supplemental Figures 1-12

Supplemental Tables 1-7

Supplemental Video 1-3

